# pcnaDeep: A Fast and Robust Single-Cell Tracking Method Using Deep-Learning Mediated Cell Cycle Profiling

**DOI:** 10.1101/2021.09.19.460933

**Authors:** Yifan Gui, Shuangshuang Xie, Yanan Wang, Ping Wang, Renzhi Yao, Xukai Gao, Yutian Dong, Gaoang Wang, Kuan Yoow Chan

## Abstract

**Motivation:** Computational methods that track single-cells and quantify fluorescent biosensors in time-lapse microscopy images have revolutionised our approach in studying the molecular control of cellular decisions. One barrier that limits the adoption of single-cell analysis in biomedical research is the lack of efficient methods to robustly track single-cells over cell division events.

**Results:** Here, we developed an application that automatically tracks and assigns mother-daughter relationships of single-cells. By incorporating cell cycle information from a well-established fluorescent cell cycle reporter, we associate mitosis relationships enabling high fidelity long-term single-cell tracking. This was achieved by integrating a deep-learning based fluorescent PCNA signal instance segmentation module with a cell tracking and cell cycle resolving pipeline. The application offers a user-friendly interface and extensible APIs for customized cell cycle analysis and manual correction for various imaging configurations.

**Availability and Implementation:** pcnaDeep is an open-source Python application under the Apache 2.0 licence. The source code, documentation and tutorials are available at https://github.com/chan-labsite/PCNAdeep.

**Supplementary Information:** Supplementary data are available online.

## Introduction

Image analysis at the single-cell level is a powerful tool to study cell-to-cell differences in a population, allowing the elucidation of cellular patterns invisible to population-averaged measurements (Skylaki *et al*., 2016). By combining multidimensional microscopic imaging with fluorescent reporters, it is now possible to study signalling processes that lead to cell fate decisions in detail. Among the most widely used fluorescent reporters are the human DNA helicase B (HDHB) (Gu *et al*., 2004), FUCCI (Sakaue-Sawano *et al*., 2008) and proliferative cell nuclear antigen (PCNA) (Zerjatke *et al*., 2017). The combination of these fluorescent reporters in various configurations have been used to reveal many new insights on the nature of cell cycle control (Barr *et al*., 2017; Spencer *et al*., 2013; Feringa *et al*., 2016; Min *et al*., 2020; Cappell *et al*., 2016; Cura Costa *et al*., 2021). A major challenge often faced when analysing these large-volume image datasets is the lack of a robust, efficient, and accurate cell detection & tracking solution (Skylaki *et al*., 2016).

Here, we developed an application that incorporates the cell cycle phase information to facilitate the long-term tracking of single cells across cell division events. We utilized a widely available fluorescently-tagged PCNA, as an all-in-one reporter, to label cells and extract cell cycle information through a deep neural network Mask R-CNN (He *et al*., 2017). This allowed us to accurately detect cell division events and construct cell lineages in a context-aware manner. We encapsulated the above functionalities with a user-friendly interface for model training, evaluation and manual correction into a Python package named pcnaDeep. To our knowledge, pcnaDeep is the first fully automated image-to-quantification solution for linage tracing and interrogating cell cycle dynamics using a single fluorescence marker.

## Methods

We worked on confocal microscopy images of RPE1 and MCF10A cells expressing endogenous mScarlet-PCNA (**Supplementary Methods 1**). The distinct changes in the PCNA fluorescence pattern during cell cycle progression faithfully report the cell cycle phases (Zerjatke *et al*., 2017). To simultaneously identify cell objects and cell cycle phase information, we performed instance segmentation by using a Mask R-CNN neural network. The model was trained on 728 images containing 34,137 cell instances, manually annotated with morphological labels that represent the various stages of cell cycle progression (**Supplementary Methods 2**). We used both PCNA fluorescence and brightfield (BF) images as the model input because the rounding of mitotic cells is easily identified on BF images (**Supplementary Experiment 1**). Cell objects were linked through the TrackPy package (Allan *et al*., 2019) which generates non-bifurcate tracks in a feature space.

To construct cell lineages with mitosis events, pcnaDeep first identifies mitosis events in TrackPy outputs. We developed a Greedy Phase Searching (GPS) algorithm (**Supplementary Methods 3**) to detect targeted phases in a noisy background. Tracks with detected mitosis phase are broken into mother and daughter tracks at the frame of maximum velocity, as an approximation of cytokinesis. These separated tracks are put into a pool of potential mother-daughter tracks together with orphan tracks that have M phase labels at the terminal. Mother-daughter relationships are joined from this pool using a spatial-temporal thresholding algorithm. For filtered tracks, a score is calculated based on the linear sum of spatial and temporal penalties, which generates a cost matrix for one-mother-to-two-daughter assignments. The cost matrix is solved by Hungary algorithm (Kuhn, 1955) to find valid mother-daughter pairs.

To quantify other cell cycle phase transitions and durations, tracks are analyzed individually. An assumption was made that the cell cycle transition of individual tracks should proceed in the following order: M-G1-S-G2-M. We applied GPS to search for S phase under the background of G1/G2 labels. The rest G1/G2 labels are resolved based on their temporal locations relative to S and M phases. An overview summarising the analytical workflow of pcnaDeep is shown in **Figure. 1**. A detailed description of the software architecture is available in the attached supplementary information (**Supplementary Methods 3**).

**Figure 1.**
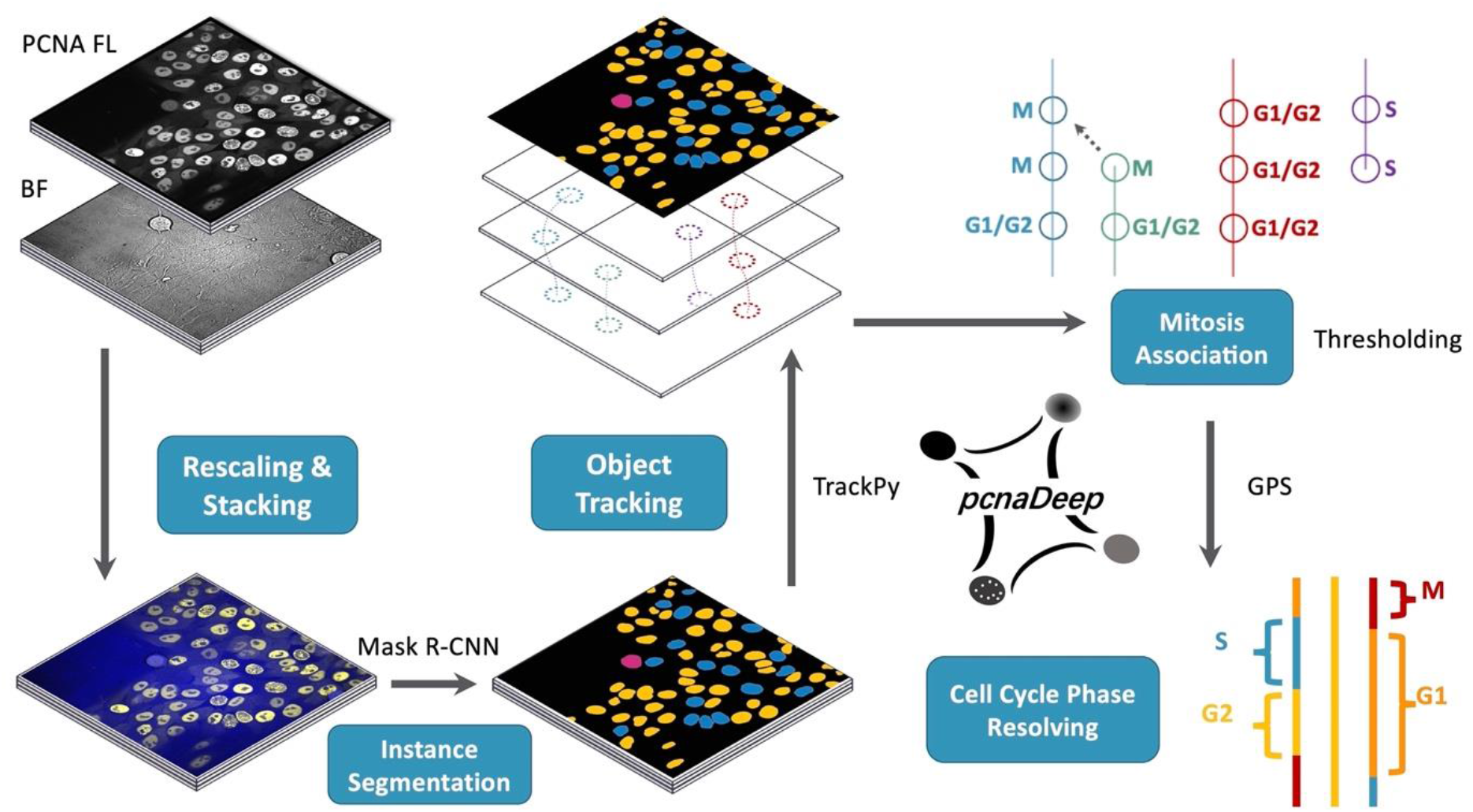
Overview of the stages in pcnaDeep main application. Mask-RCNN identifies cell nucleus instances and their morphological labels on PCNA fluorescence (PCNA FL) plus bright-field (BF) composite videos. TrackPy links objects into single tracks. Mitosis relationships are predicted for candidate tracks through thresholding, which link tracks into lineages. Cell cycle phases are resolved for each lineage using the GPS algorithm.

## Results

We annotated the instance segmentation and cell cycle ground truth of six time-lapse videos to evaluate pcnaDeep **(Supplementary Methods 4, Supplementary Experiments 2~4)**. The weighted-average F1 score (wmF1) across cell cycle phase transitions or complete phase categories were adopted as major evaluation metrics. **(1)** To benchmark against TrackMate (Tinevez *et al*., 2017), we substituted tracking and mitosis association steps with TrackMate linear assignment problem (LAP) tracker. Compared with TrackMate, pcnaDeep shows much higher accuracy in determining cell cycle phase transitions (pcnaDeep wmF1: 0.93±0.02; TrackMate: 0.64±0.11; mean±s.d., 6 videos) and cell cycle phase detection (pcnaDeep: 0.90±0.03; TrackMate: 0.39±0.12). **(2)** A grid search of thresholds for mitosis association showed low parameter sensitivity, with optimal configurations resulting in a mitosis phase detection accuracy of 0.94, recall of 0.81 and F1 of 0.87. **(3)** Down-sampling videos by half did not influence the performance with accordingly scaled parameters. **(4)** The frame error of cell cycle phase transition and duration was similar to a panel of human labellers.

The efficient Mask R-CNN model and downstream designs enable pcnaDeep to process hundreds of image frames with high cell density within minutes (**Supplementary Experiments 5**). By analyzing the cell cycle feature, mitosis tracks can be accurately associated, demonstrating the usefulness of this cellular context in cell tracking. Moreover, the output of pcnaDeep is importable into Fiji ImageJ (Schindelin *et al*., 2012) for visualization and quantification. If required, manual correction can be done through a command-line interface.

## Conclusion

We present pcnaDeep as an application that automatically segments the nucleus and tracks the cells in a context-aware manner, providing a flexible output mask useful for quantifying cellular dynamics of molecules in long-term live imaging experiments. pcnaDeep significantly saves time by removing the requirement for human manual tracking and annotation of single-cell tracks that take ~3 days/100 cells to several minutes. The accuracy of pcnaDeep is comparable to human annotations and do not suffer from human experimental bias, making this an attractive cell cycle profiling approach. Importantly, fluorescently tagged PCNA have been successfully introduced into a variety of biological systems using transient and stable transfection methods (Leonhardt *et al*., 2000; Kisielewska *et al*., 2005; Zerjatke *et al*., 2017; Icha *et al*., 2016; Barr *et al*., 2017). This makes pcnaDeep a powerful and flexible tool to accurately track and generate nuclear masks of cells in a variety of experimental settings that require single-cell linage tracing.

## Acknowledgements

We are grateful to Jonathon Pines and Oxana Nashchekina for their kind guidance in using PCNA as a fluorescent cell cycle reporter. We would like to thank Shengyu Hao and members of the Gaoang Wang lab for their fruitful discussions. We would also like to express our appreciation to Wanlu Liu for critically reading our manuscript and for her helpful suggestions.

