## Supplementary material for "pcnaDeep: A Fast and Robust Single-Cell Tracking Method Using Deep-Learning Mediated Cell Cycle Profiling": online

#### Supplementary Materials

### Supplementary Methods

#### 1 Tissue Culture Experiments

##### 1.1 Cell Culture

hTERT-RPE1 (CRL-4000; ATCC) cells were grown in Dulbecco's Modified Eagle Medium/Nutrient Mixture F-12 Ham (Sigma-Aldrich) medium supplemented with GlutaMAX (Thermo Fisher), 10% FBS (Gibco), 0.348% sodium bicarbonate, penicillin (100 U/ml), streptomycin (100 µg/ml) at 37°C with 5% $CO_2$. The female human non-transformed breast epithelial cell line MCF10A (CRL-10317; ATCC) was maintained in Dulbecco's Modified Eagle Medium/Nutrient Mixture F-12 Ham (Sigma-Aldrich) supplemented with 5% horse serum, 20 ng/ml EGF, 0.5 mg/ml Hydrocortisone, 100ng/ml Cholera toxin, 10 µg/ml Insulin, GlutaMAX (Thermo Fisher), 100U/ml penicillin and 100mg/ml streptomycin at 37°C with 5% CO2 according to established protocols (Debnath *et al.*, 2003). Cells were passaged every 2-3 d with trypsin (Sigma-Aldrich) and not allowed to reach confluency except where intentionally arrested. All cell lines have been tested for mycoplasma contamination.

##### 1.2 Generation of PCNA-reporter cell lines

hTERT-RPE1 and MCF10A mScarlet-PCNA cells were generated as described by the marker-free co-selection method (Agudelo *et al.*, 2017). eSpCas9 (Addgene plasmid #86613) and ATP1A1 repair template sequence used in the co-selection method, is a gift from Iain Hagan's lab. PCNA guide RNA sequences (GCGCGCCTCGAACATGGTGG) were selected using CRISPOR (Concordet and Haeussler, 2018) and cloned into the eSpCas9 plasmid using the Bbs1 restriction site. The PCNA repair template with mNeonGreen fused to the N-terminus of the PCNA coding sequence was synthesized by GenScript. Vector maps and sequences are available upon request.

##### 1.3 Live-cell time-lapse microscopy

For time-lapse microscopy, 1.0 x $10^4$ hTERT-RPE1 cells and 3.0 x $10^4$ MCF10A cells were seeded on µ-Slide 8 Well chamber slides (ibidi), two days before being imaged under the microscope. Slides were placed in an Okolab stage incubator (OKO) at 37°C with 5% $CO_2$, and 80% humidity. A Nikon Ti2 inverted fluorescence microscope equipped with a spinning-disk system (CSU-X1; Yokogawa) and operated with the MetaMorph software was used to image the cells. The cells were observed under a 60x plan apo objective (NA 1.2) and images were captured using a Photometrics Prime 95B camera. The configuration used by the mCherry filters were 562/40 nm EX, 624/40 nm EM. Imaging conditions varied according to the cell line. For hTERT-RPE1-mScarlet-PCNA, a 4 × 1µm z-stack was captured in both DIC and mCherry at 5min intervals for up to 24h. For imaging of MCF10A-mScarlet-PCNA, a 5 × 2µm z-stack was captured in both DIC and mCherry at 5min intervals for up to 24h. Images were in the size of 1200*1200 square pixels (1.83 µm/pixel).

### 2 Dataset Preparation

#### 2.1 Segmentation Ground Truth Labelling with Cell Cycle Morphological Annotation

The instance segmentation task in pcnaDeep is to identify the cell nuclei during interphase and the whole cell during mitosis, along with its cell cycle stage. As described (Zerjatke *et al.*, 2017) and illustrated in **Fig. S1**, fluorescent PCNA signal forms morphologically distinct foci in the cell nucleus during S phase. In G1 and G2, PCNA signal distributes evenly in the cell nucleus (**Fig. S1a**). Although the PCNA intensity level is higher in G2 compared to G1, this subtle difference may be masked by the unstable illumination during time-lapse imaging (**Fig. S1b**). Therefore, both G1/G2 stages are grouped as a single classification during the segmentation stage. During mitosis, the rounding up of the cells occasionally resulted in the loss of the PCNA signal (**Fig. S1b**). Therefore, bright-field images were used to identify the mitotic state and to provide a clear cell boundary (**Fig. S1a**). To address cells that just completed cytokinesis which was morphologically unique (as the nuclear envelope has reformed but the PCNA signal is still cytoplasmic), we introduced an "emerging" phase (E) classification other than G1/G2, S and M. We refer to the above semi-cell cycle phase classifications as morphological labels.

The segmentation ground truth on pixel-level was initially labelled through conventional thresholding and manual correction. Then we adopt Mask R-CNN (He *et al.*, 2017), one of the state-of-the-art instance segmentation approaches, to facilitate annotation. After training the Mask R-CNN model with a considerable number of images, we generated the ground truth by correcting the model prediction output. All manual work was done through the VGG Image Annotator (Dutta and Zisserman, 2019) by a group of well-trained human labellers and was reviewed at least two times to ensure accuracy. Specific criteria for morphological labelling are described in **Table S1**.

#### 2.2 Cell Cycle Ground Truth Labelling

Cell tracking, mother-daughter assignments and cell cycle phase ground truths are embodied in the tracked object table. For each video, a tracked object table was obtained by using segmentation ground truth as the model input. A command-line interface was used to correct inaccurate object identities and cell cycle classifications within the table. Tracks shorter than 10 frames (50 minutes) were excluded from the ground truth. Arrested G1 or G2 cells that could not be accurately classified due to the presence of uneven illumination and the lack of phase transition information were excluded from the analysis. The command-line correction interface is released as a part of the application.

#### 2.3 Datasets

728 training images with mCherry (PCNA) and bright-field channels were used for training the Mask R-CNN model. 122 testing images were used for evaluating the instance segmentation performance. 34,138 object instances in the training images and 6,231 in the testing images had been labelled (**Table S2**). For evaluating the entire application, 6 videos taken at 0.2 frames per minute, lasting ~300 frames on average had been labelled. The testing videos cover a wide range of cell densities and mitosis event counts (**Table S3**). For training the support vector machine (SVM) (Platt and others, 1999) mitosis association classifier, we used three videos taken with lower frequency at 0.05 frame per minute (**Table S4**).

### 3 Software Architecture

#### 3.1 Mask R-CNN Training

The Mask R-CNN model was trained on a baseline (mask_rcnn_R_50_FPN_3x) provided by Facebook's Detectron2 framework (Wu *et al.*, 2019). This configuration has a 50-layer residual network (He *et al.*, 2015) + feature pyramid network (Lin *et al.*, 2017) backbone and had been pre-trained on the Microsoft COCO instance segmentation dataset (Lin *et al.*, 2017). By adapting Detectron2's training schedule, the baseline model was trained for additional 2800 iterations with 16 images per batch on our 728-composite-image training set (~62 epochs, Method 2.3). The base learning rate was set to 0.1 with a warmup period lasting 400 iterations followed by a series of 10-fold learning rate decay at iteration 800, 1400, 1800 and 2200. Composite images were generated by stacking two same PCNA fluorescent images with one bright-field image into the RGB format. Composite images were scaled to 1% saturation and then augmented with the following approaches to reduce their intercorrelations, especially for those sampled from videos: random cropping by 0.8*0.8 times; random flipping; random saturation by 0.5~1.5 times and random rotation for either 0, 90 or 270 degrees. Other training configurations followed Detectron2 v0.4 defaults. Model training was carried out on an NVIDIA GeForce RTX-3090 GPU under CUDA 11.0, PyTorch 1.7.1 and Ubuntu 18.04.

#### 3.2 Instance Segmentation

To perform instance segmentation on videos (image stacks), the intensity level of each frame in the PCNA fluorescent and bright-field stacks is scaled to 1% saturation by default. After rescaling, composite images are generated in the same way as in Mask R-CNN training. From each detected object in the composite image, essential fields for later stages are extracted. Objects smaller than 800 pixel$^2$ or locates within 10 pixels of the image border are filtered out. As for overlapping detections, objects with higher classification confidence scores overwrite those with lower scores on the output mask. Object classifications are smoothed by averaging confidence scores through a 5-frame-long sliding window. Objects classified as E are treated the same as G1/G2 in the above steps.

#### 3.3 Multiple Object Tracking

Object tracking is performed by the `link` method implemented in the TrackPy package (Allan *et al.*, 2019) that utilizes the Crocker-Grier algorithm (Crocker and Grier, 1996) for tracking objects in an n-dimensional feature space. pcnaDeep uses temporal & spatial locations, as well as the mean bright-field signal intensity of the objects as the input features. In the `link` method, the maximum movement of the object in the feature space between consecutive frames (`search_range`) is set to 80 pixels by default; the memory of objects for filling gapped tracks (`memory`) is set to 10 frames by default; `adaptive_stop` is set to 0.4*`search_range`. TrackPy does not recognize cell division events, which is addressed in a dedicated stage below.

#### 3.4 Mitosis Detection and Association

In a mother-daughter mitotic relationship, TrackPy links the mother cell with one of the closer daughter cells in the feature space. The mitotic events within tracks are detected using the Greedy Phase Searching (GPS) algorithm. Within detected mitosis periods, mother and daughter objects are separated at the time point when the object moving speed maximizes, assuming cells move the most at cytokinesis. Multiple rounds of mitosis are allowed since such separation process is recursively performed until no more mitosis can be detected. This separation process generates candidate mitotic mothers. Additional candidate mothers are defined as those having at least one mitosis (M) classification during their disappearance period (10 frames by default). Likewise, candidate mitotic daughters are defined as those tracks containing at least one M or emerging (E) classification during their appearance period (10 frames by default), together with daughters separated from their mothers. Afterwards, the spatial ($dD$) and temporal ($dT$) differences (**Table S5**) of every pair of mother and daughter tracks in the candidate pool are extracted. Two modes generating the probability score of a candidate mitotic relationship have been implemented. The thresholding mode is used by default:

1) The thresholding mode (TRH, default): Spatial and temporal features are first filtered by user-specified thresholds (default spatial: 120 pixels; temporal: 10 frames). Scores are calculated for the remaining from the weighted sum. A balanced weight $W_d = 0.5$ is adopted by default.
$$Score = 1 - W_d \cdot dD - (1 - W_d) \cdot dT$$
2) The supervised SVM mode (SVM): An SVM model with linear kernel taking the same shaped input as in TRH ($dD$ and $dT$) is fitted on the training data of three ground truth videos (**Table S4**). Positive instances are extracted from valid mother-daughter pairs, and the negative instances are extracted from all invalid pairs formed by tracks involved in positive pairs. Data are pre-processed by a quantile-based scaler (sklearn.preprocessing.RobustScaler) to ensure the robustness of handling the outliers (Pedregosa *et al.*, 2012). The C parameter has been optimized through 5-fold cross-validation grid search on the labelled training set ($C = 13$). During testing, the training set is augmented with the run-time positive instances, i.e., mother and daughter tracks separated from a single track, aiming to capture more testing set-specific features. The probability score of a valid relationship is directly assigned as the SVM output confidence.
$$Score = SVM(\ dD, dT)$$

To normalize different cell movements across different cell lines and image sampling frequency, the spatial ($dD$) and temporal ($dT$) features are calculated as in **Table S5**. The spatial (pixel unit) and temporal (frame unit) thresholds are normalized by $\bar{R} + \overline{dD}$ and $SAMPLE\_FREQ$ at runtime respectively.

Based on the assumption that one mother cell can only divide into at most two daughter cells, a cost matrix is constructed with rows representing mothers and columns representing daughters. Mothers in the rows are duplicated to allow a one-to-two assignment. Each cost in the matrix is the opposite of the $0 \sim 1$ probability score of the corresponding pair of mother and daughter tracks. For mother-daughter pairs initially separated from single tracks, the cost is set to $-1$. The cost matrix is optimized via the Hungary algorithm (Kuhn, 1955) to find the configuration with the least sum. Mitosis relationships are established for matched mother-daughter pairs in the optimized configuration.

#### 3.5 Cell Cycle Phase Resolving

After mitosis detection, the mitosis phase duration can be directly calculated. Since tracks are already separated by mitosis events, the rest phases should follow the G1-S-G2 order. Complete S phases are detected by the GPS algorithm and G1/G2 phases are inferred from their temporal location relative to S. Incomplete phases are partially resolved by putting a ‘>’ notation before their duration value in the output table. Tracks are classified as arrested if no phase transition is detected in the above steps. For arrested G1/G2 tracks, the user can specify an intensity threshold (background subtracted) to distinguish G1 from G2.

#### 3.6 Greedy Phase Searching Algorithm (GPS)

The algorithm aims to identify a period with consistent target cell cycle phase classifications out of the noisy background (**Fig. S2**). By accumulating confidence scores of targeted or background classification confidence scores, a period is accepted or rejected according to an over-threshold target score $S \geq T_{TG}$ or background score $P \geq Th_o$, respectively. The target threshold $Th_{TG}$ and the background threshold $Th_O$ are set to 5 by default. The background and target score are reset to zero every time $S$ exceeds the target threshold, conferring the algorithm with generalisability over different target period lengths (assuming longer tracks have more faulty classifications). The $fe$ (free ending) optional flag lifts the criteria at the track terminal for identifying mitosis mother tracks with unstable detections. Major variables are explained in **Table S6**.

**Algorithm** Greedy Phase Searching

```
Function Search(L, TG, C, Th_TG, Th_o, e, fe)
    Find all i such that L_i = TG
    I_1, I_2, ... , I_N ⟵ i_k, i_{k+1}, ... , k ≥ e
    S, P ⟵ 0
    t_entry ⟵ I_1
    Found ⟵ False
    For t ⟵ 1 ... N do
        S ⟵ S + C_{I_{t+1}}
        P ⟵ P + Σ_{1+I_t}^{-1+I_{t+1}} C
        If S ≥ Th_TG
            Found ⟵ True
            If P < Th_O
                S, P ⟵ 0
        If P ≥ Th_O
            If Found
                t_exit ⟵ I_t
                Break Loop
            Else
                S, P ⟵ 0
                t_entry ⟵ I_{t+1}
    If P < Th_O AND ( Found OR
                ( fe AND I_N − t_entry + 1 ≥ Th_TG AND t_entry ≠ I_N))
        t_exit ⟵ I_N
        Found ⟵ True
    Return t_entry, t_exit if Found, else NA
```

#### 3.7 False-proof Measures

Four false-proof measures have been adopted to inform the user that the inference result may not be accurate. **1**) For mitotic daughters associated but do not have any mitosis classification, the mitosis exit cannot be precisely measured. This is indicated in one of the model outputs (cell cycle phase table). **2**) For lineages that do not contain a detected S period between two consecutive mitoses, the cell cycle information is not resolved for the entire lineages, which is indicated during runtime. **3**) Invalid transitions like S-M are indicated during runtime. **4**) Tracks showing >20% phase difference after resolving may be highly noisy and do not follow the phase transition order. These tracks are indicated during runtime. With the above measures, the user can easily spot and correct faulty tracks.

### 4 Evaluation Methods

#### 4.1 Instance Segmentation Evaluation

Segmentation result was scored via Cell Cycle Challenge SEG measurements as a reference (Ulman *et al.*, 2017). The SEG score is based on calculating mean inversion over union (mIoU) in the cell masks. We used a 122-image testing set for evaluation (**Table S2**). The segmentation performance was first benchmarked against conventional Otsu thresholding methods through Fiji ImageJ 1.53c (Schindelin *et al.*, 2012) using the following macro:

```
run("Enhance Contrast...", "saturated=1");
run("Gaussian Blur...", "sigma=2 stack");
setAutoThreshold("Otsu dark");
setOption("BlackBackground", false);
run("Convert to Mask", "method=Otsu background=Dark calculate");
run("Open", "stack");
run("Fill Holes", "stack");
run("Analyze Particles...", "size=800-Infinity pixel show=Masks stack");
run("Watershed", "stack");
run("Invert LUT").
```

For benchmarking against Cellpose v0.6 (Stringer *et al.*, 2021), we used a pre-trained 3-model average with an estimated diameter of 80 pixels. For benchmarking against Deepcell v0.9.1 (Moen *et al.*, 2019), we used the NuclearSegmentation application in Deepcell and scaled images to 1% saturation before input. The size parameter micron per pixel was set to *image_mpp = 0.183*. Unmentioned parameters were kept the same as the corresponding defaults.

#### 4.2 Cell Cycle Profiling Evaluation

To evaluate the entire model, we first matched objects in the ground truth to testing output via least-sum assignment (Hungary algorithm) of the centroid distance matrix in each frame. Assigned object pairs that had centroid distances larger than 40 square pixels were rejected. Objects in the testing output were matched to the ground truth in the same way.

To evaluate the accuracy of phase transition detection, all phase transition periods were extracted from the ground truth tracks, each expanding a 30-frame window at maximum. Every ground truth transition period was matched to the testing data if all the objects within the period were mapped to testing objects belonging to a unique track for non-mitotic transitions, or a unique mother-daughter pair of tracks for mitotic transitions. Similarly, all phase transition periods in testing data were matched to the ground truth. These two mutual matchings were merged based on shared track identities. After merging, transitions detected in ground truth periods but not in testing were defined as false negative (FN), detected in testing but not in ground truth as false positive (FP) and the remaining true positive (TP). Likewise, complete cell cycle phases were matched based on shared object and track identities to evaluate the accuracy of complete phase detection.

For each transition and phase category, the Recall, Precision and F1 scores were calculated as the following:

$$Recall = \frac{TP}{TP + FN}; \quad Precision = \frac{TP}{TP + FP}; \quad F1 = 2 \cdot \frac{Precision \cdot Recall}{Precision + Recall}$$

We calculated the weighted mean of F1 scores across different cell cycle transition/phase categories in each testing video as our major objectives. We regarded the absolute frame error of all TP detections as another objective.

#### 4.3 Comparison with TrackMate

The cell tracking and mitosis association were compared to the linear assignment problem (LAP) tracker implemented in Fiji TrackMate plugin (Tinevez *et al.*, 2017) using the parameter setting: *max dist = 40; max gap = 10; max gap dist = 40; max splitting dist = 120*. The detection output was imported into TrackMate using the CSVImporter plugin. To resolve the cell cycle phases from TrackMate output, all bifurcate tracks were identified and split into mother-daughter pairs. If no mitosis classification could be found at the track termini, the mitosis classification was assigned to the beginning/end of the daughter/mother tracks. All other resolving parameters & steps were kept the same as pcnaDeep default.

#### 4.4 Comparison with Human Labellers

Three labellers (L1~3) were assigned with annotating the MCF10A-01 video independently based on their empirical criteria of determining the cell cycle progression. Labellers were given the tracking ground truth, and they could decide either using bright field, PCNA fluorescent, or both images to complete the task. Labellers were asked to recover as many cell cycle phase transitions and durations as possible. Detection accuracy and frame errors compared with the ground truth were calculated.

#### 4.5 Testing on Half-sampled Data

Raw images and ground truth object tables were sampled once every two frames to generate half-sampled datasets. In the processed ground truth, no parent track lost mitosis classifications. For ground truth daughter tracks losing mitosis classification after half-sampling, the mitosis exit was assigned to the first frame of the track's appearance. During testing, all time-related configurations in pcnaDeep were divided by two, including frame window for classification smoothing, sampling frequency, temporal gap during tracking, appearance thresholds in GPS, mitosis search range, frame threshold for associating mitotic tracks and minimum track length to resolve. The maximum object movement in tracking and distance threshold for mitosis association were scaled by 1.5 times. The phase transition window length for evaluation (**Method 4.2**) was set to 15. Only cell cycle transitions & phases detected in both half-sampled and original datasets were considered when comparing the absolute frame errors.

### Supplementary Results

#### 1 Instance Segmentation

The distinct appearances of PCNA fluorescent (FL) signal during cell cycle progression faithfully report the cell cycle phases. Such a feature, however, poses the challenge of generating cell segmentation masks through conventional thresholding approaches, as PCNA does not distribute evenly during S phase and does not stain the nucleolus. Using mIoU-based SEG measurement as proposed in the Cell Tracking Challenge (Tinevez *et al.*, 2017) on a 122-image testing set, we observed SEG=0.49 when using conventional Otsu thresholding on PCNA fluorescent images. The pixel-level threshold does not capture object-level information, therefore cannot resolve the unevenly distributed PCNA signal well. From the deep-learning perspective, the available pre-trained neural networks did not perform well (SEG score, Cellpose (Stringer *et al.*, 2021) 3-model average: 0.69; Deepcell (Moen *et al.*, 2019): 0.49) especially on mitotic cells, possibly because PCNA fluorescent images are not represented in common cell segmentation training sets. Regarding the uniqueness of our task in resolving the cell cycle phase, we carried out transfer learning of a Mask R-CNN neural network baseline (He *et al.*, 2017) under the Detectron2 framework (Wu *et al.*, 2019) to perform cell cycle instance segmentation (**Method 4.1**). Owing to the clear cell rounding feature during mitosis in the bright-field (BF) images, training using FL plus BF images outperformed FL images only as for the mitosis classification (**Table S7**), as evaluated by the Microsoft COCO (Lin *et al.*, 2014) average precision (AP). The FL plus BF trained model reached an SEG score of 0.92. Hence, we adopted both FL and BF images as essential inputs for the main application. An example of the segmentation performance is shown in **Figure S3**.

#### 2 Accuracy of Cell Cycle Phase Transition and Duration

pcnaDeep was applied to six testing videos covering a wide range of cell density and mitosis events (**Table. S3**) using default parameters. For comparison, we substituted the tracking and mitosis association modules in pcnaDeep with TrackMate linear assignment problem (LAP) tracker (**Method 4.2, 4.3**). The performance of the entire application was first evaluated from the cell cycle phase transition perspective. By aligning phase transition windows in the detected tracks with the ground truth, we classified each window as either TP, FP, FN, or biologically invalid transitions. The precision, recall and F1 scores were calculated accordingly. In sharp contrast to TrackMate which tracks objects unaware of the cell cycle phase information, pcnaDeep outputs much fewer faulty detections (**Fig. S4a, b**) and shows higher accuracy in mitosis-related categories (G2-M, M-G1, **Fig. S4c**). The weighted-average F1 score across transition categories is significantly higher for pcnaDeep (0.93±0.02, mean±s.d.) compared with TrackMate (0.64±0.11). As for the Cell Cycle Challenge TRA tracking evaluation (**Table. S8**), although pcnaDeep showed slightly higher TRA scores in all six videos, the effect is subtle, probably because TRA penalizes mitotic error to a less extent. As for the entire cell cycle phase detection, pcnaDeep outperformed TrackMate in aspects of fewer mitotic FP and FN cases (**Fig. S5**), which resulted in a similarly large contrast of the weighted mean F1 score (pcnaDeep: 0.90±0.03; TrackMate: 0.39±0.12).

Most of the frame errors of TP phase transitions in pcnaDeep output of the six testing videos was kept within ±2 frames (**Fig. S6a**). The phase duration errors of non-mitotic phases were mostly limited within ±5% of the ground truth averages (**Fig. S6b**). We then compared the performance of pcnaDeep with three humans on labelling the MCF10A-01 dataset (one of the six testing videos, **Method 4.4**). pcnaDeep displayed F1 scores that were comparable to human labellers in recognising cell cycle phase transitions (**Fig. S6c**). As for the complete cell cycle phases, pcnaDeep detected G1 and G2 at the accuracies similar to human labellers, although it performed slightly worse in recognizing M and S phases (**Fig. S6d**). The absolute frame error of cell cycle transition had a large variance among human labellers, while pcnaDeep displayed an error range similar to human labellers except for G2-M (**Fig. S6e**). pcnaDeep sometimes reported mitosis transitions late, which is likely caused by the late detection of the mitotic state. Likewise, the frame error of phase durations was highly variable (**Fig. S6f**) among humans while pcnaDeep again exhibited an error range comparable to humans. Therefore we concluded that the accuracy of pcnaDeep is comparable to human labelling. We note that tracks having large transition frame errors may not be present in the phase frame error plots as they may not contain complete phases and were therefore excluded from the analysis.

#### 3 Thresholding and SVM in Mitosis Association

In pcnaDeep, the mitotic mother-daughter relationship is established by resolving a one-mother-to-two-daughter assignment matrix. The cost of each candidate pair of mother and daughter is generated from calculating the linear penalization score on threshold-gated distance & frame gaps by default. The cost can also be assigned as the output of an SVM classifier processing on the same spatial-temporal features, in a supervised manner. To compare these two methods, we applied a parameter space of distance (40~200 pixels) and frame (5~20 frames) thresholds to the detection output and plotted the precision-recall curve (P-R curve) of mitosis phase detection in the six testing videos. Similarly, an array of confidence thresholds was set to filter the SVM output score to plot the P-R curve (**Fig. S7**). Unlike common P-R curves directly plotted from a binary classifier, in both conditions, the curve is bounded within a small zone, which is resulted from the subsequent mother-daughter pair assignment step. The performance of the thresholding approach (**Fig. S7a**) is not sensitive to the distance threshold once it is set close to the cell size (~80 pixels in our datasets). The method is also not sensitive to the frame threshold around half of the estimated 15-frame mitosis length. On the other hand, the SVM model (**Fig. S7b**) requires low (<0.1) confidence thresholds to reach the recall (>0.75) comparable with the thresholding approach. Moreover, the SVM never outperformed thresholding in the present experiments. Hence, the current SVM model fitted on the three low-frequency training videos (**Table. S4**) is not generalizable.

#### 4 Generalization Across Imaging Frequency

When profiling the cell cycle, the trade-off between phototoxicity and sampling frequency leads to various choices of time-lapse imaging configurations. To test whether pcnaDeep is generalizable in the aspect of imaging frequency, we down-sampled testing videos by half and generated the corresponding ground truth. When applying pcnaDeep with scaled parameter settings (**Method 4.5**), the application showed similar accuracy in detecting both cell cycle phase transition (**Fig. S8a**) and duration (**Fig. S8b**). The weighted mean F1 score on half-sampled samples (transition: 0.92±0.03; phase: 0.90±0.03) resembles those of the original samples (transition: 0.93±0.02, phase: 0.90±0.03). We next compared the frame error of phase transition and duration using cases detected in both situations (95.1% of original; 97.2% of half-sampled, **Fig. S9**). There is no difference in the absolute error distributions when contrasting the transition (**Fig. S9a**) or duration (**Fig. S9b**) output with the ground truth of the corresponding frequency, suggesting that scaling the parameter in different frequencies does not influence the model performance. However, the error is amplified when comparing the half-sampled result with the original ground truth, especially for phase transition detection, indicating that sampling frequency may have an impact on the accuracy of profiling the cell cycle phase.

#### 5 Runtime

The runtime (**Table. S3**) of six videos under pcnaDeep default configurations was tested on two platforms: 1) A high-performance server with Intel Xeon Gold 6139M 2.3 GHz CPU, NVIDIA GeForce RTX-3090 GPU, 128 GB RAM, Ubuntu 18.04 64-bit operation system; 2) A regular workstation with AMD Ryzen 7 2700X 3.7 GHz CPU, NVIDIA GeForce RTX-2060 GPU, 32 GB RAM, Windows 10 Pro 64-bit operation system. The runtime increased with higher cell density and longer time frames but was kept within several minutes.

### Supplementary Figures

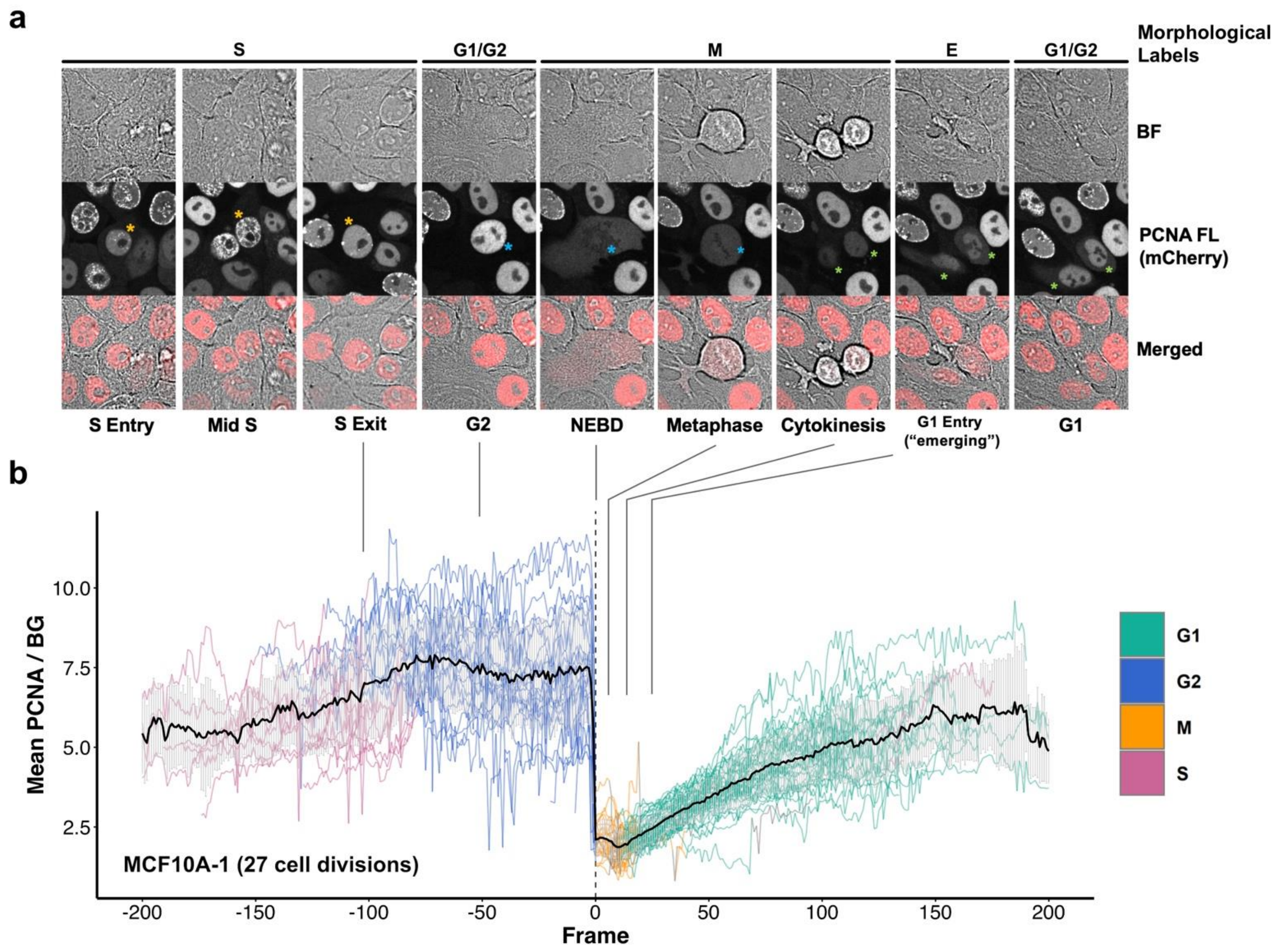

**Supplementary Figure S1. Morphological labelling example of cell cycle phases. (a)** Representative confocal images of MCF10A cells in different cell cycle phases and the corresponding morphological labels. Stars of the same colour refer to the same cell. Cells labelled with green stars are daughters of the cell labelled with a blue star. BF: bright-field; FL: fluorescent. **(b)** PCNA signal intensity dynamics in 27 lineages involving cell division of a representative dataset MCF10A-01 sampled at 0.2 frames per minute. Lineages were aligned at mitosis entry (nuclear envelope breakdown, NEBD). For each labelled cell mask, the PCNA signal intensity was calculated as the averaged pixel intensity within the object mask and normalized with the background intensity (BG) which was calculated by averaging all background pixels within the 4-time dilated bounding box area of the object. The average signal across cell tracks is shown as the bold line in the middle (error bar: s.d.). Arbitrary temporal locations of panels in (a) are linked to (b).

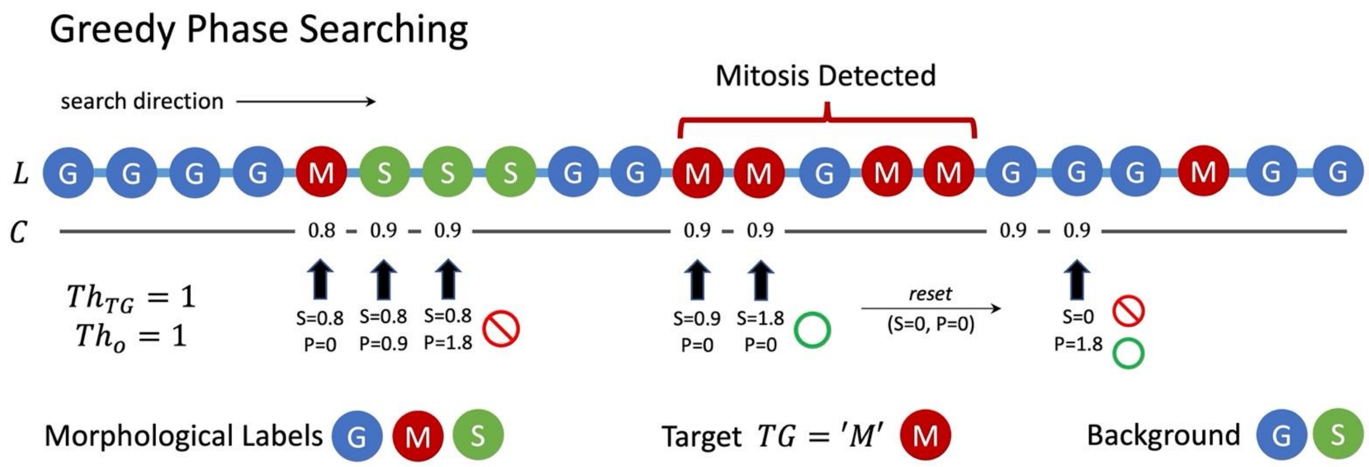

**Supplementary Figure S2. Schematic of the Greedy Phase Searching Algorithm being applied to detect mitosis.** For searching mitosis along the sequence of morphological labels $L$ and the embedding confidence score $C$, the accumulative target $S$ and background $P$ confidence scores are calculated. The mitosis phase is detected or rejected according to the over-threshold target ($S \geq Th_{TG}$) or background ($P \geq Th_O$). $S$ and $P$ are reset every time $S \geq Th_{TG}$ happens.

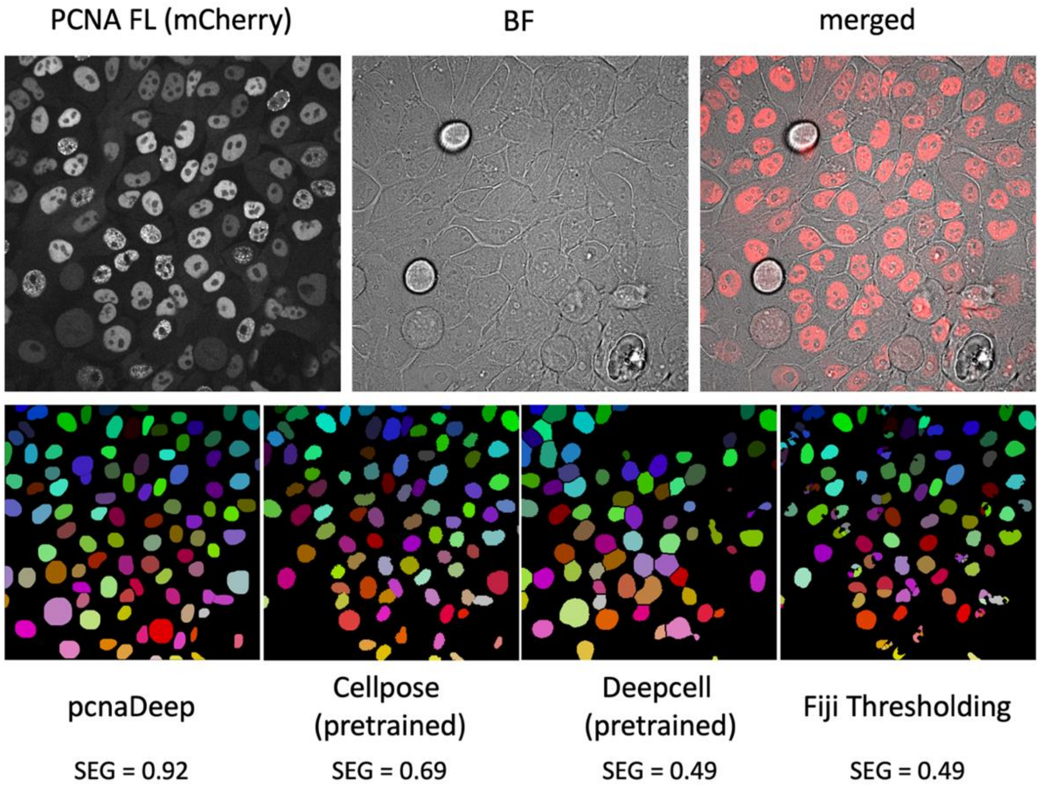

**Supplementary Figure S3. Example of the segmentation performance on an MCF10A image slice.** The segmentation mask was stained by pseudocolour. pcnaDeep takes both fluorescent (FL) and bright-field (BF) channels as the input while the others take FL only.

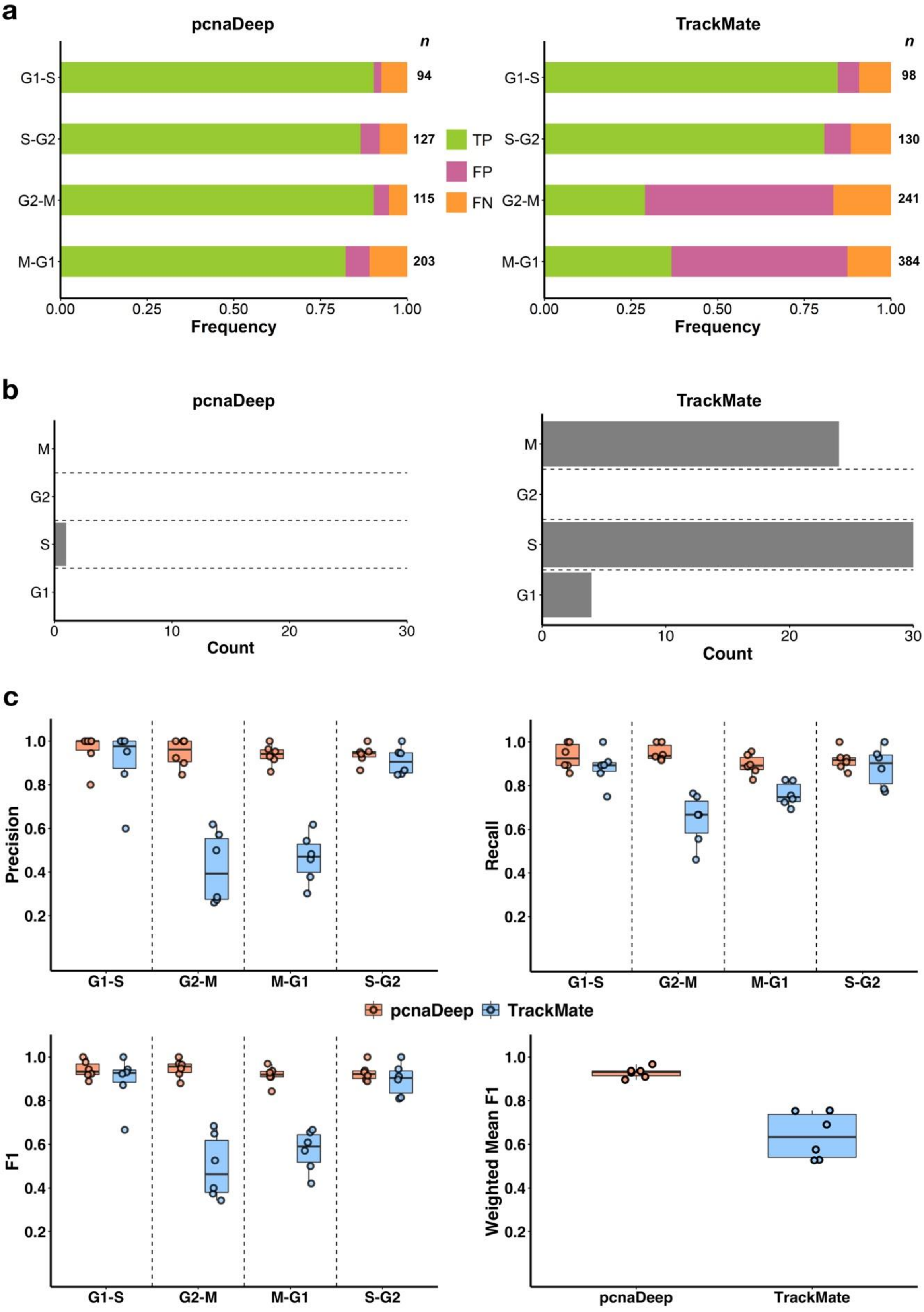

**Supplementary Figure S4. Cell cycle phase transition detection performance of pcnaDeep and TrackMate. (a)** Frequency of detected TP/FP/FN cell cycle phase transitions. **(b)** Quantification of biologically invalid transitions in the output of all testing videos. Invalid transitions in (b) were grouped by the cell cycle phase of the transition entry (y-axis). **(c)** Detection evaluation matrices calculated from data in (a). The weighted F1 score was averaged across the four-phase transition categories and weighted by the number of transitions in the output. Each point in the plots corresponds to one video (six videos in total).

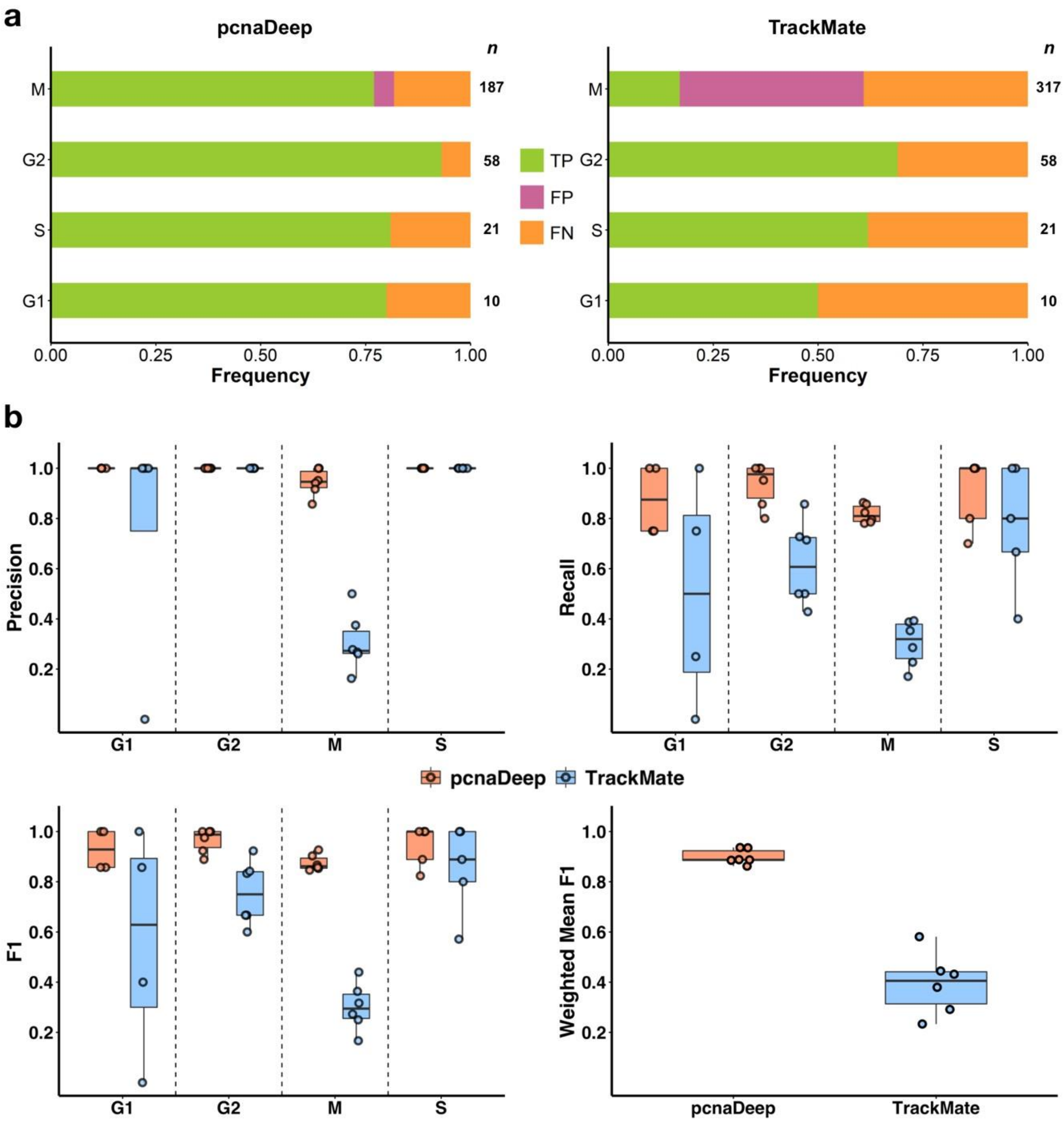

**Supplementary Figure S5. Cell cycle phase duration detection performance of pcnaDeep and TrackMate. (a)** Frequency of detected TP/FP/FN cell cycle phase in the output of all testing videos. **(b)** Detection evaluation matrices calculated from data in (a). The weighted F1 scores were averaged across the four-phase categories and weighted by the number of detected phases in the output. Each point in the plots corresponds to one video (six videos in total). Two videos without G1 and one video without S phase in the ground truth are not plotted in the corresponding column in (b).

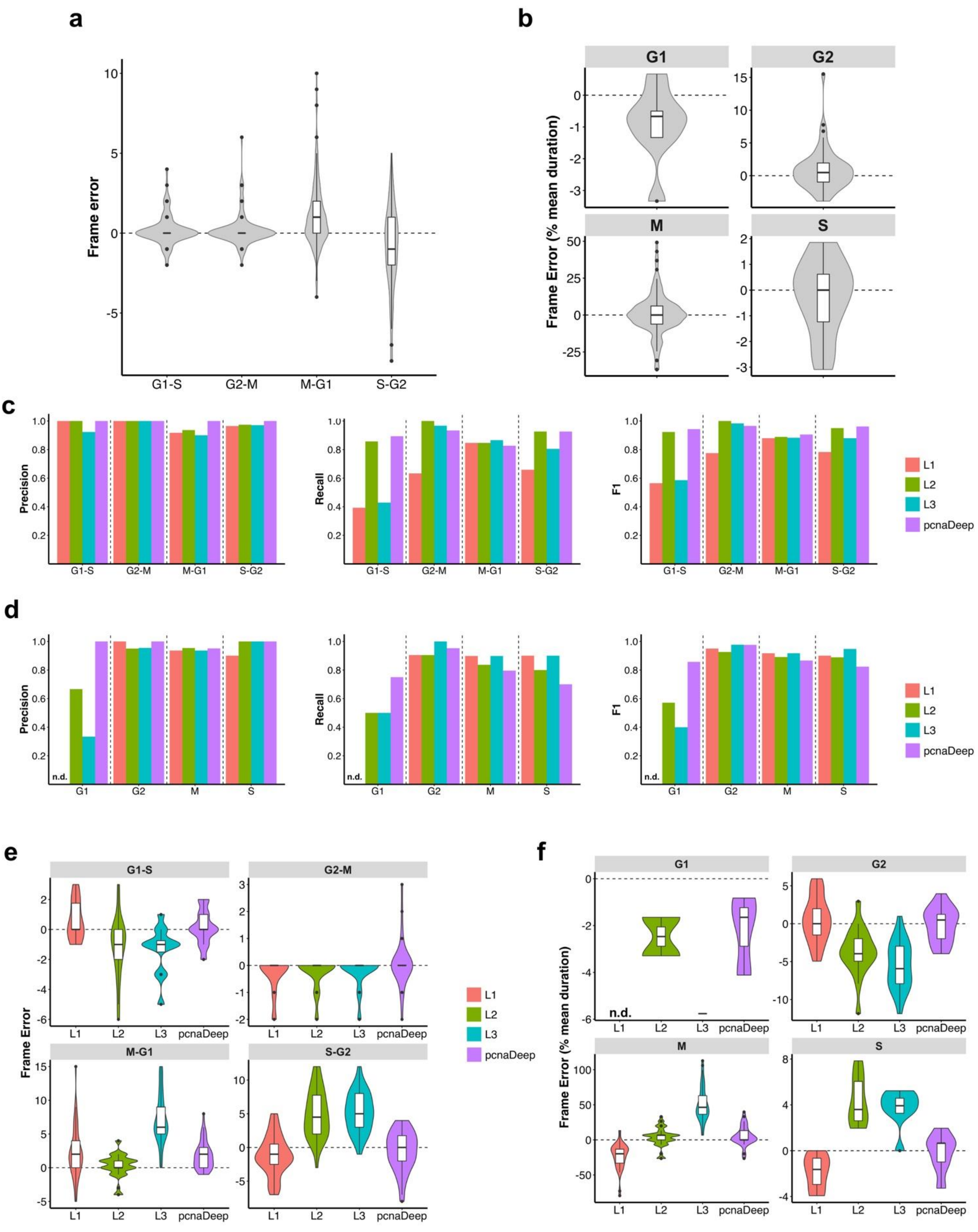

**Supplementary Figure S6. Frame error of pcnaDeep in recognizing the cell cycle progression and comparison with human labellers. (a)** Absolute frame error of pcnaDeep in recognising TP cell cycle phase transitions from six testing videos. **(b)** Frame error of pcnaDeep in determining TP phases relative to averaged duration in the ground truth from six testing videos. (c)~(f): Comparison between human labellers (L1, L2 and L3) and pcnaDeep on analysing dataset MCF10A-01. **(c)** Evaluation matrices for the detection of cell cycle phase transitions. **(d)** Evaluation matrices for the detection of the duration of the cell cycle phases. **(e)** Absolute frame error of TP transitions in (c). **(f)** Frame error of TP phases in (d) normalized by averaged duration in the ground truth. (0.2 frames per minute; n.d. not determined; L1 did not label any G1; sample size in (a) and (b): G1-S: 85, G2-M: 104, M-G1: 167, S-G2: 110, G1: 8, S: 17, G2: 54, M: 144; sample size in the MCF10A-01 ground truth in (c)~(f): G1-S: 28, G2-M: 30, M-G1: 52, S-G2: 41, G1: 4, S: 10, G2: 21, M: 49).

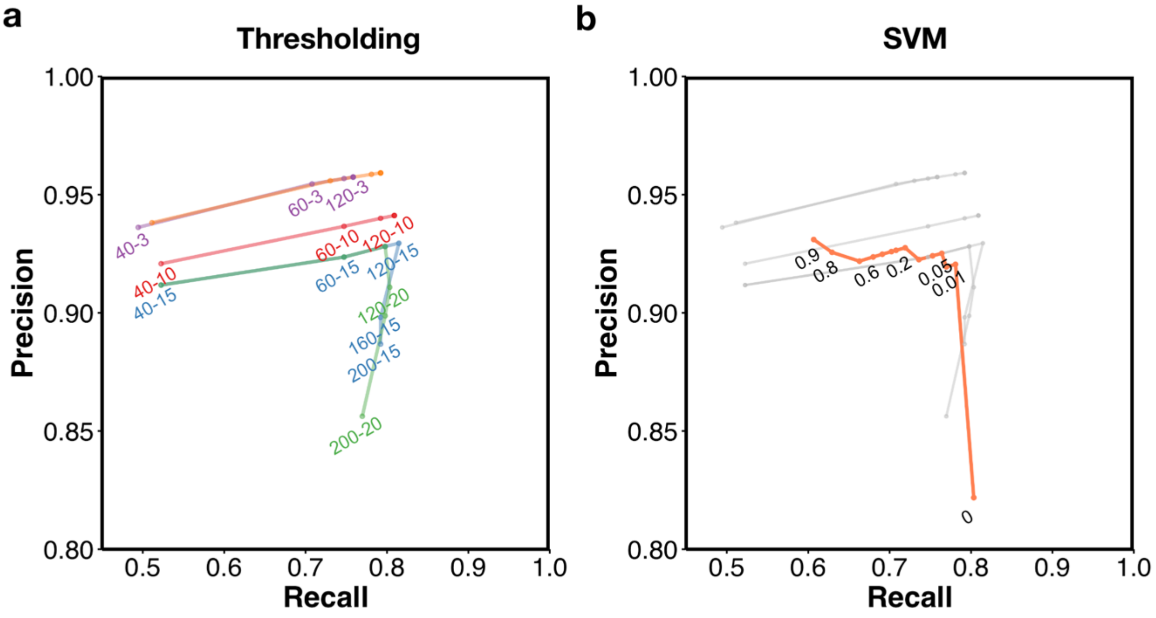

**Supplementary Figure S7. Precision-Recall curve of mitosis phase detection. (a)** Thresholding approach. "60-10" means distance threshold = 60 pixels and frame threshold = 10 frames. Configurations with the same frame threshold are linked together. **(b)** SVM approach. The number refers to the threshold imposed on SVM confidence output. The SVM P-R curve is superimposed on the thresholding P-R curves for comparison. Note each point in (a) and (b) corresponds to one configuration.

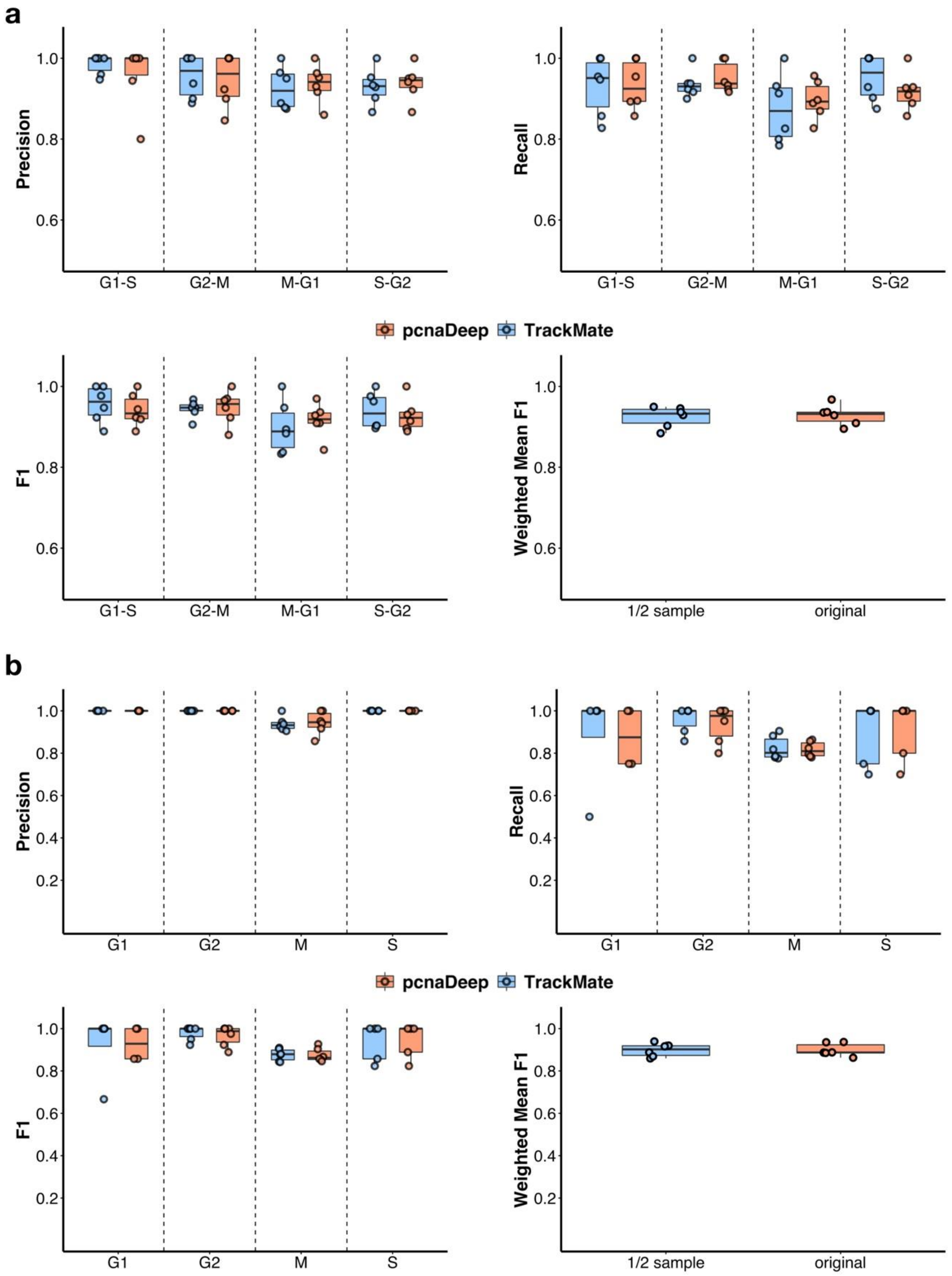

**Supplementary Figure S8. Detection performance on half-sampled videos. (a)** Detection evaluation matrices for cell cycle phase transition. **(b)** Detection evaluation matrices for complete cell cycle phases. The detection performance on both (a) and (b) compares either original or half-sampled (1/2 sample) ground truth under either condition.

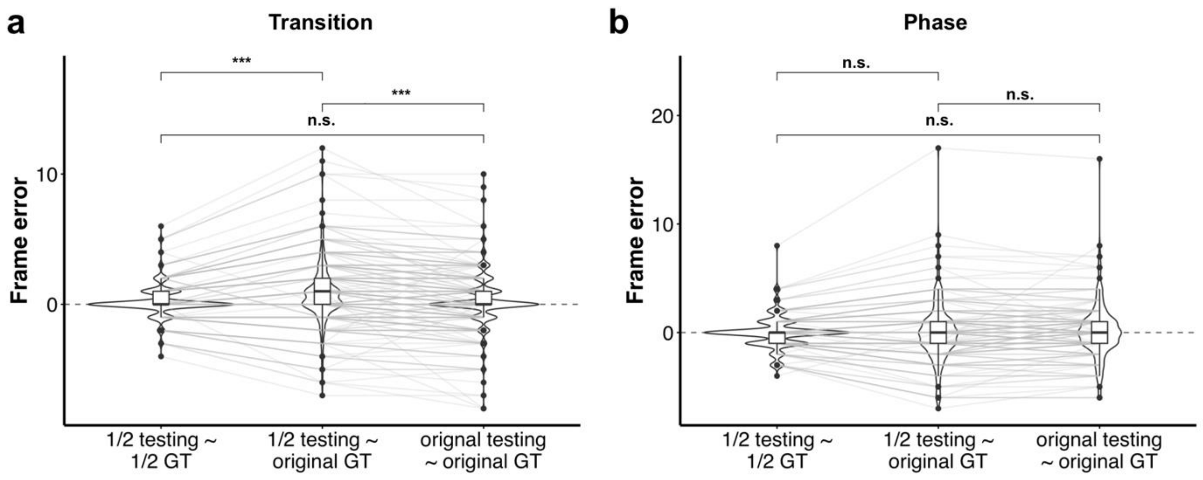

**Supplementary Figure S9. Frame accuracy of cell cycle phase transition and duration on half-sampled videos. (a)** Absolute frame error of true positive cell cycle phase transition (n=448). **(b)** Absolute frame error of total cell cycle phase duration (n=207) in the testing output of all six videos were either compared with the half-sampled or the original ground truth, as indicated on the x-axis. All categories of cell cycle transition or phase were pooled together. Paired *t*-test (n.s. no significance; *** p<0.001).

### Supplementary Tables

| Classification | Criteria |
|---|---|
| **S** | At least three visible PCNA foci were found in the cell nucleus. |
| **M** | After nuclear envelope breakdown, with clear cell rounding features (dark boundary in the bright-field channel images), until the formation of clear nucleus boundary. |
| **E** | After the formation of the nuclear boundary, with relatively lower contrast between cytoplasm and nucleus PCNA signal compared with surrounding G1/G2 cells. |
| **G1/G2** | The rest cases. |

**Supplementary Table S1. Morphological annotation criteria in the segmentation stage.**

| Training | Images | G1/G2 | S | M | E | Total |
|---|---|---|---|---|---|---|
| **MCF10A** | 417 | 19438 | 5515 | 1498 | 828 | 27729 |
| **RPE1** | 311 | 4426 | 2179 | 169 | 84 | 6408 |
| **Total** | 728 | 23864 | 7694 | 1667 | 912 | 34137 |
| **Testing** | **Images** | **G1/G2** | **S** | **M** | **E** | **Total** |
| **MCF10A** | 65 | 3059 | 1466 | 175 | 71 | 4771 |
| **RPE1** | 57 | 1047 | 363 | 32 | 18 | 1460 |
| **Total** | 122 | 4106 | 1829 | 207 | 89 | 6231 |

**Supplementary Table S2. Counting of images and instances used for training & testing the Mask R-CNN model.**

| Video ID | Frame length | Frequency (FPM) | Average cells per frame | Mitosis events | Runtime[1] (sec) | Runtime[2] (sec) |
|---|---|---|---|---|---|---|
| **MCF10A-01** | 265 | 0.2 | 86.49 | 49 | 210 | 322 |
| **MCF10A-02** | 135 | 0.2 | 58.67 | 22 | 87 | 127 |
| **MCF10A-03** | 186 | 0.2 | 88.30 | 41 | 159 | 214 |
| **RPE1-01** | 343 | 0.2 | 31.67 | 17 | 165 | 309 |
| **RPE1-02** | 433 | 0.2 | 27.54 | 29 | 194 | 263 |
| **RPE1-03** | 410 | 0.2 | 28.87 | 21 | 226 | 272 |

**Supplementary Table S3. Six time-lapse videos used for evaluating the application and the runtime.** A mitosis event refers to one mother-daughter cell pair, i.e., one cell division corresponds to at most two mitosis events. Only one pair was tracked when the other daughter cell moves out of the image field. FPM: frame per minute. Runtime on a high-performance server (1) and a regular workstation (2) are shown.

| Video ID | Frame length | Frequency (FPM) | Average cells per frame | Mitosis events |
|---|---|---|---|---|
| **MCF10A-04** | 66 | 0.05 | 87.38 | 43 |
| **MCF10A-05** | 66 | 0.05 | 80.56 | 41 |
| **RPE1-04** | 65 | 0.05 | 27.48 | 18 |

**Supplementary Table S4. Three time-lapse videos used for training the SVM mitosis classifier.**

| Feature | Formula | Normalization factor |
|---|---|---|
| Distance difference $dD$ | $\sqrt{(x_m - x_d)^2 + (y_m - y_d)^2}$ | $\bar{R} + \lvert dT \rvert \cdot \overline{dD}$ |
| Frame difference $dT$ | $t_d - t_m$ | $SAMPLE\_FREQ$ |

**Supplementary Table S5. Calculation of spatial and temporal features for mitosis association.** $x_m, y_m, x_d, y_d$: mother and daughter cell location; $t_m$: frame of mother cell's disappearance; $t_d$: frame of daughter cell's appearance; $\bar{R}$: average radius of all objects, estimated as averaged short and long axis length of the fitted ovals; $\overline{dD}$: average movement across frames of all objects; $SAMPLE\_FREQ$: sampling frequency (frame per minute), which is a user-specified metadata constant.

| Variable | Description |
|---|---|
| $L$ | A list of detected cell cycle phase classifications. |
| $TG$ | Target cell cycle phase to search. |
| $C$ | Confidence of the classifications, same shape as $L$. |
| $Th_{TG}$ | Accumulated confidence threshold for detecting the target phase. |
| $Th_O$ | Accumulated confidence threshold for detecting other phases. |
| $e$ | Exclude the first $e$ detections of the target. |
| $fe$ | Flag to lift requirements for detecting target phase at the track end. |
| $S$ | Accumulative target confidence score. |
| $P$ | Accumulative background confidence score. |

**Supplementary Table S6. Glossary of variables involved in the Greedy Phase Searching algorithm.**

| | AP50 | AP75 | APm | AP-G1/G2 | AP-S | AP-M | AP-E |
|---|---|---|---|---|---|---|---|
| FL + BF | 84.25 | 76.89 | 70.29 | 85.83 | 88.93 | **77.35** | 26.75 |
| FL only | 77.91 | 70.83 | 62.23 | 85.56 | 88.81 | 48.96 | 28.78 |

**Supplementary Table S7. Evaluation of the Mask R-CNN Training.** The network was trained on either fluorescent (FL) plus bright-field (BF) images or FL only. The evaluation matrix is the Microsoft COCO average precision (AP). AP50/75: AP under IoU 0.5/0.75; APm: median object AP; AP-G1/G2, AP-S, AP-M, AP-E: AP of corresponding morphological classifications.

| CTC Matrix | MCF10A-01 | MCF10A-02 | MCF10A-03 | RPE1-01 | RPE1-02 | RPE1-03 |
|---|---|---|---|---|---|---|
| SEG | 0.896 | 0.923 | 0.911 | 0.917 | 0.920 | 0.910 |
| pcnaDeep TRA | 0.967 | 0.986 | 0.974 | 0.984 | 0.981 | 0.967 |
| TrackMate TRA | 0.963 | 0.985 | 0.972 | 0.983 | 0.980 | 0.967 |

**Supplementary Table S8. Evaluation with CTC matrices.** SEG for segmentation measurement and TRA for tracking measurement on the final output. In TrackMate experiments, only the tracking and mitosis association stages of pcnaDeep were replaced with the TrackMate LAP tracker.
